# Review of layperson screening tools and model for a holistic mental health screener in lower and middle income countries

**DOI:** 10.1101/763045

**Authors:** Aderibigbe Oluwakemi Olanike, Christopher M Perlman

## Abstract

**Background:** The needs of people diagnosed with Mental Neurological and Substance-Use (MNS) conditions are complex including interactions physical, social, medical and environmental factors. Treatment requires a multidisciplinary approach including health and social services at different levels of care. However, due to inadequate assessment, services and scarcity of human resource for mental health, treatment of persons diagnosed with MNS conditions in many LMICs is mainly facility-based pharmacotherapy with minimal non-pharmacology treatments and social support services. In low resource settings, gaps in human resource capacity may be met using layperson health workers. A layperson health working is one without formal mental health training and may be equivalent to community health worker (CHW) or less cadre in primary health care system.

**Objectives:** This study reviewed layperson mental health screening tools for use in supporting mental health in developing countries, including the content and psychometric properties of the tools. Based on this review this study proposes recommendations for the design and effective use of layperson mental health screening tools based on the Five Pillars of global mental health.

**Methods:** A systematic review was used to identify and examine the use of mental health screening tools among laypersons supporting community-based mental health programs. PubMed, Scopus, CINAHL and PsychInfo databases were reviewed using a comprehensive list of keywords and MESH terms that included mental health, screening tools, lay-person, lower and middle income countries. Articles were included if they describe mental health screening tools used by laypersons for screening, delivery or monitoring of MNS conditions in community-based program in LMICs. Diagnostic tools were not included in this study. Trained research interviewers or research assistants were not considered as lay health workers for this study.

**Results:** There were eleven studies retained after 633 were screened. Twelve tools were identified covering specific disorders (E.g. alcohol and substance use, subcortical dementia associated with HIV/AIDS, PTSD) or common mental disorders (mainly depression and anxiety). These tools have been tested in LMICs including South Africa, Zimbabwe, Haiti, Malaysia, Pakistan, India, Ethiopia and Brazil. The included studies show that simple screening tools can enhance the value of laypersons and better support their roles in providing community-based mental health support. However, most of the layperson MH screening tools used in LMICs do not provide comprehensive information that can inform integrated comprehensive treatment planning and understanding of the broader mental health needs of the community.

**Conclusion:** Developing a layperson screening tools is vital for integrated community-based mental health intervention. This study proposed a holistic framework which considers the relationship between individual’s physical, mental and spiritual aspect of mental health, interpersonal as well as broader contextual determinants (community, policy and different level of the health system) that can be consulted for developing or selecting a layperson mental health screening instrument. More research are needed to evaluate the practical application of this framework.

## Introduction

### Mental Health System and MNS Conditions in LMICs

The prevalence of mental, neurological and substance use (MNS) disorders in lower-middle income countries is about two in ten persons(1). It is estimated that 76%-85% of persons affected by MNS disorders in LMICs lack access to mental health services for prevention and treatment resulting in a huge treatment gap(1,2). Mental disorders, if untreated, can cause significant impairment and disability resulting in emotional and economic burden on individuals, their families, caregivers and the society at large. Mental, neurologic and substance use disorders are one of the ten leading causes of disability globally, accounting for over 11% of the global burden of disease (GBD) measured by disability adjusted life years (DALY) and 28% of GBD measured by years lived with disability (YLD) in 2016(3). The magnitude of disability caused by MNS conditions results from early onset of the illness, failure to seek help or delay in initiating treatment. These are in part due to lack of knowledge about mental disorder and available treatment, unavailability of care and barrier caused by stigma(2).

Mental, neurological and substance use disorders also affect the overall quality of life of people with MNS disorders and their caregivers. In some jurisdictions, people with mental illness are denied basic rights and are faced with numerous societal barriers especially those arising from stigma and discrimination(4). This in turn can affect their ability to fully participate as members of their societies. Their inability to work constitute economic burden on their families, particularly due to the costs of formal and informal care. The emotional impact includes distress associated care, stigma and lives lost to suicide. The quality of life of people with MNS disorder is worse in LMICs where they are usually neglected in poor living conditions and with no access to quality health care. Also, government expenditure earmarked for mental health in LMIC is not proportionate to the contribution of mental health to disease burden. It was estimated that an average 0.5% of total health expenditures are allocated for mental health in LMICs. This imposes an enormous challenge on the health care system(5).

The needs of persons with mental health conditions are complex, including interactions among physical, social, cultural, medical and environmental factors. Therefore the treatment of people diagnosed with MNS disorders ideally requires multidisciplinary approaches including health and social services at different levels of care. However, this is not always the case in low-resource settings like most LMICs where treatment of persons diagnosed with MNS conditions is mainly facility-based pharmacotherapy with minimal non-pharmacology treatments and social support services.

The lack of integrated assessment tools could explain the gap in their treatment plan because it is impossible to get information about what is not assessed for. And lack of information about other factors contributing to the general wellbeing of persons with MNS conditions results in treatment gap. There is need for integrated and multi-sectoral approach to the assessment of need of people with MNS conditions in order to provide them with holistic treatment.

In 2001, the World Health Organization (WHO) published a landmark report on mental health, with the goal of increasing public awareness on the burden of mental disorders and removing barriers that are creating treatment gaps for those that need care. In order to reduce the treatment gap of MNS conditions the WHO proposed recommendations that can be adapted by every country to support people living with MNS conditions(6). Key interventions recommended for improving MH in LMICS include empowering people with MNS disorders and their families to provide support to each other, training non specialist health workers to deliver psychological treatments, integrating economic intervention into mental health care, use of computer-assisted, self-guided psychological therapies, delivering school – based interventions for childhood disorders and providing integrated care for people with mental disorders(5). These strategies have been classified broadly as the integration of mental health services into primary healthcare, expansion of human resources or capacity for mental health through task sharing and training of non-specialist and various innovation to engage lay person in self-care and informal community care in order to enhance access, reduce cost and reduce stigma(7). These strategies create the opportunity to expand the integration of mental health into existing health care system, strengthen human resources, improve delivery of services and care reaching more persons with MNS conditions.

### Laypersons Support of Mental Health in LMICS

Engagement of non-professionals, or lay persons, in the screening and delivery of mental health may be a promising mechanism to improve support for persons with MNS conditions. Non-specialist mental health workers have been classified as health professionals/workers that may have received general mental health training but are not specifically trained as mental health professionals (e.g. doctors, nurses, para-professionals and nonprofessional lay providers)(8,9). Non-specialist mental health workers may also include professionals that are not involved in health care directly but play important roles in mental health promotion and detection, such as teachers and community level workers, parents, traditional healers, village elders, community based volunteers and peers(8,9). Evidence shows that lay persons can be engaged in promotion and primary prevention, identification and detection, treatment, care and rehabilitation of MNS disorders(10).

Layperson services are commonly used to provide mental health and psychosocial support in contexts where there is scarcity of human resources. There are several examples in which lay community workers or volunteers have been trained to deliver high quality mental health and psychosocial support interventions under the supervision and guidance of trained professionals(8,9,11). In these instances, lay workers also provided basic psychosocial support, group-based counselling, symptom management and referral for specialist psychological support with the aim of reducing distress, improving psychological and psychosocial functioning and improving coping mechanism of individuals and their community. Services provided by these community volunteers also include individual screening/evaluation using different tools(8,9,11). There are many existing system opportunities for the integration of non-health or non-mental health workers into mental health workforce. Lay workers can conduct mental health screening and referral to PHC or hospitals, mental health screening may be integrated into social activities in the community (E.g. schools, social clubs, and religious centers) and other community based organizations (CBO) activities like health outreaches and emergency response can be integrated as support systems. At the crux of these opportunities is the need for a common mechanism, or tool, to support lay workers/volunteers to be able to properly identify and support the needs of those in their communities.

### Mental Health Screening Tools in LMICs

Screening tools can be used to provide succinct information about the needs and resources of persons with MNS condition and the community in which they live. Tools can also be used for large scale epidemiological research to determine the mental health of a community or population. Content within screening tools can also be used for training and creating community awareness to reach more people with MNS conditions and improve access to care through community-based activities and outreach.

Mental health instruments can be classified based on the purpose they serve and the MNS condition they are used to identify. Typically, mental health screening tools are used for screening, diagnosis or treatment monitoring/evaluation(12), while comprehensive assessment tools help to gather detailed information that is needed for accurate diagnosis and treatment plan that meets the individual need of the patient.(13). Diagnostic tools provide information that are useful for specialist/clinicians to determine the nature and or cause of the presenting complaints in order to make a diagnosis according to DSM classification. Screening tools are different from diagnostic tools in that they provide information used to identify those at risk of MNS disorder and that might need further evaluation by a specialist. Treatment monitoring and evaluation tools are used to track changes in symptoms and functioning to determine the effectiveness of the treatment/intervention. While some instruments fall categorically under one of these classifications, some could actually be used for two or three of these purposes. It is explicit that diagnostic tools are used by mental health specialist or health workers with adequate mental health training, but some screening tools and treatment monitoring tools can be used by trained laypersons.

A variety of mental health screening tools have been developed and applied to detect MNS conditions in LMICS(14). Some studies have pooled validation studies of mental screening tools in general and some for specific mental conditions in particular populations (12,14). A number of these tools require administration by trained professionals while others can be administered by persons with no formal mental health training (14). While tool inventories exist, there is limited evaluation of the feasibility and effectiveness of layperson mental health screening tools, particularly for community-based programs in LMICs. The purpose of this study was to review layperson mental health screening tools for use in supporting mental health in developing countries, including the content and psychometric properties of the tools. Based on this review this study will propose recommendations for the design and effective use of layperson mental health screening tools based on the Five Pillars of global mental health.

## Methods

This study uses a systematic review to identify tools and examine their use among laypersons supporting community-based mental health programs. The study was conducted in accordance with the PRISMA recommendations for systematic reviews(15). There was no study protocol published in advance of conducting this review.

### Search Strategy

The following keywords were used to conduct the literature search in a systematic manner: mental health, screening tools, lay-person, lower and middle income countries. Combinations of search terms, as shown in Fig 1, were used to identify manuscripts from PubMed, Scopus, CINAHL and psychInfo databases. These databases search was restricted to journals published between 2008 and 5^th^ June 2018.

**Fig 1:**
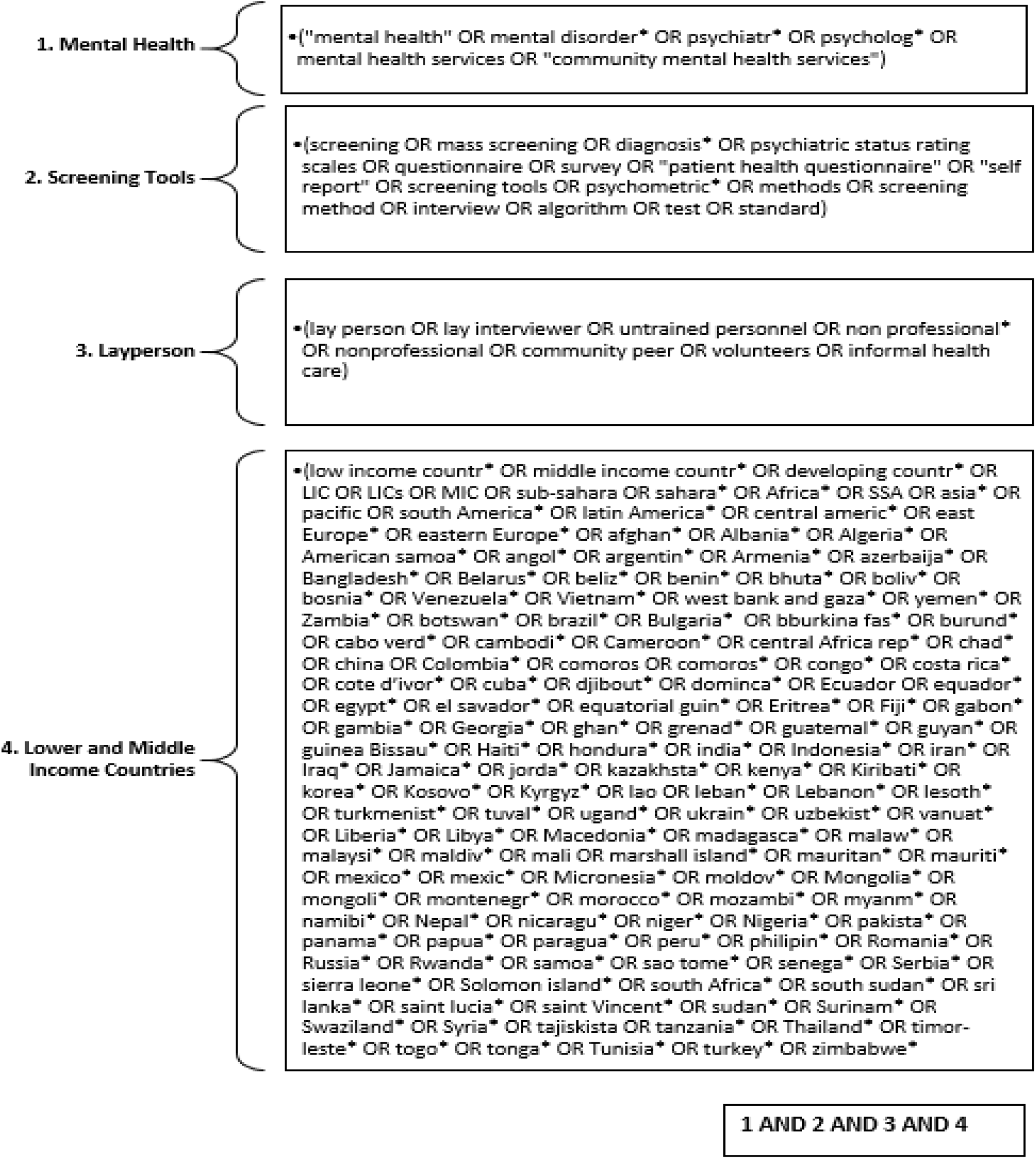
Search terms used to identify manuscripts from the databases.

### Inclusion and Exclusion Criteria

Articles were included if they describe mental health tools for use by laypersons for screening, delivery or monitoring of MNS conditions in community-based program in LMICs. Laypersons have been described above as persons that are non-mental health professionals and non-clinicians or non-health workers supporting mental health program or intervention (e.g. teachers, parents, peers, local/community workers, community health worker). Diagnostic tools were not included in this study because these are not expected to be used by laypersons without clinical training. Trained research interviewers or research assistants were also not considered as lay assessors for this study because we could not verify their professional or prior training. Studies in which lay persons were only engaged in providing community-based mental health interventions but not in the assessment or screening of the participants were also excluded from this review. Studies that did not specify who administered the tools were also excluded. Articles published before 2008 and in languages other than English were also excluded.

### Study Selection

First level title screening was done to excluded studies that were not related to mental health/mental health screening tools. Abstracts of the remaining articles were reviewed for possible inclusion. Full texts of all articles that were potentially relevant were assessed using the inclusion criteria. The reference lists of the articles included were searched for additional studies. The second author repeated 10% of the study selection at every stage in order to reduce bias caused by human error. The rate of agreement between the two reviewers was quite high and discrepancies were resolved through discussion. Authors of some of the articles were contacted for clarification about validation and definition of users of the screening tools when these are not specified in the studies.

### Quality Appraisal

All the studies that met the above inclusion criteria were included in this review irrespective of their methodological quality. This decision is based on scarcity of literature on the topic and the aim to maximize the use of available studies. Instead, methodological considerations were included as points of discussion to drive future research.

### Data Extraction

A data extraction template was developed and included the following: MNS conditions, number of items, cost, form, psychometric properties, other conditions assessed, age/study population, study setting, country of study, tool administrators, whether or not use required training and availability of multiple versions.

### Conceptual Framework for Study Analysis

The holistic policy and intervention framework (HPIF) also known as the five pillars of global mental health and addiction, provides guidance on the analysis, evaluation, and sustainability of global mental health capacity building interventions (Khenti et al, 2015). The development of this framework was a result of collaborative work between the office of transformative global health and its partners from LMICs based on practical experience, lessons learned and global best practices. For instance, programs aimed at developing international partnership, leadership training for mental health professionals, capacity building for mental health research and knowledge exchange in Sri Lanka and Sub-Saharan Africa demonstrated how capacity building can contribute to improved population mental health(16,17). The contextualization of mhGAP for primary health care in Nigeria also demonstrated the importance of considering the context and sociocultural relevance of mental health intervention in LMICs(18) The pillars of global mental health is a multilevel framework consisting of five central components which include: holistic health, cultural and socioeconomic relevance, partnerships, collaborative action-based education and learning and sustainability(19). The framework is multi-level in the sense that it examined mental health development interventions at multiple levels of healthcare system (social, political, economic, policy, community, organizational, interpersonal and individual levels)(19).

The overall objective of HPIF is to improve the health and quality of life of individuals around the world by supporting improvements in mental health care of diverse health systems. The development of this framework is underlined by a fundamental values of equity and human rights against the challenges of stigma and discrimination(19). The first pillar of HPIF emphasizes a *holistic perspective* towards health. This means that in order to successfully develop and implement a context-specific capacity building intervention, it is important to consider the interrelationships between individual, interpersonal, organizational, community, and policy levels of the health system, as well as the relationship between physical, mental and spiritual health. The second pillar emphasizes that interventions need to maintain *cultural and socioeconomic relevance*. Since cultures vary in their perspectives toward mental health conditions, knowledge and understanding of these conditions should come from the community of people for which mental health interventions are provided; this includes the opportunity to critically analyze the western perception of these conditions. Persons with mental health conditions should be important stakeholders contributing to the entire process of the mental health intervention, such as the development, design, planning, implementation, and evaluation of policies and interventions. This will enhance ownership of the project by the community and its sustainability. Therefore, the third pillar focuses on the importance of *collaboration between stakeholders*. For instance, it is essential to establish reciprocal partnership based on trust and respect with local stakeholders including community health workers, religious leaders, and local governance. This will serve as a platform for knowledge exchange, reconciliation of differences and embracing similarities in culture and values. Once a platform is established, the fourth pillar of *capacity building* can be enacted. Capacity building focuses on collaborative action-based education and learning that can be achieved through education and training of trainers. This action-oriented learning includes the identification of gaps, strengths and opportunities in the existing system, addressing the gaps by building on existing strengths and opportunities, training professionals and sharing knowledge. If successful, strategies to support the fifth pillar, *sustainability*, can ensure the long-term impact of mental health addiction interventions.

For this study the identified mental health screening tools were examined to evaluate their fit into the HPIF. To determine fit we addressed a number of questions when examining tools, such as: Is the development of the tool based on knowledge of the people of the community in which it will be used? Is the tool adapted, validated and reliable for use in that particular cultural, social and economic context? In addition to mental health, does it also assess physical and spiritual health? Does the use of the tool provide opportunity for training or knowledge acquisition on mental health? Is it sustainable in terms of availability, cost, ease of use and open to review and update?

## Results

### Study Selection

The initial database search on PubMed yielded 2,826 articles, Scopus yielded 33 articles, psychinfo yielded 991, and CINAHL yielded 134 articles. After removing duplicates a total of 1,953 articles were identified. After first level title screening, the abstracts of the remaining 633 articles were screened with 589 articles excluded because the study either was not conducted in LMICs, involved the use of diagnostic screening tool(s), or the screening tool(s) was not administered by trained professionals or research interviewers with formal training. The full text of the remaining 44 articles were further screened and their references scanned to identify the final 11 articles reviewed for this study.

One systematic review was identified focusing on the review of mental health screening tools that are validated for use in LMICs. This review did not focus specifically on tools validated for lay person use. However, three articles cited in the review that involved laypersons in the validation studies were included in this study. The eleven articles that were reviewed for this study were those which describe use of mental health screening tools by lay persons for screening, delivery or monitoring of MNS conditions in a community-based mental health programs in LMICs(14,20,29,21–28).

See PRISMA Flowchart (Fig 2) below

**Fig 2:**
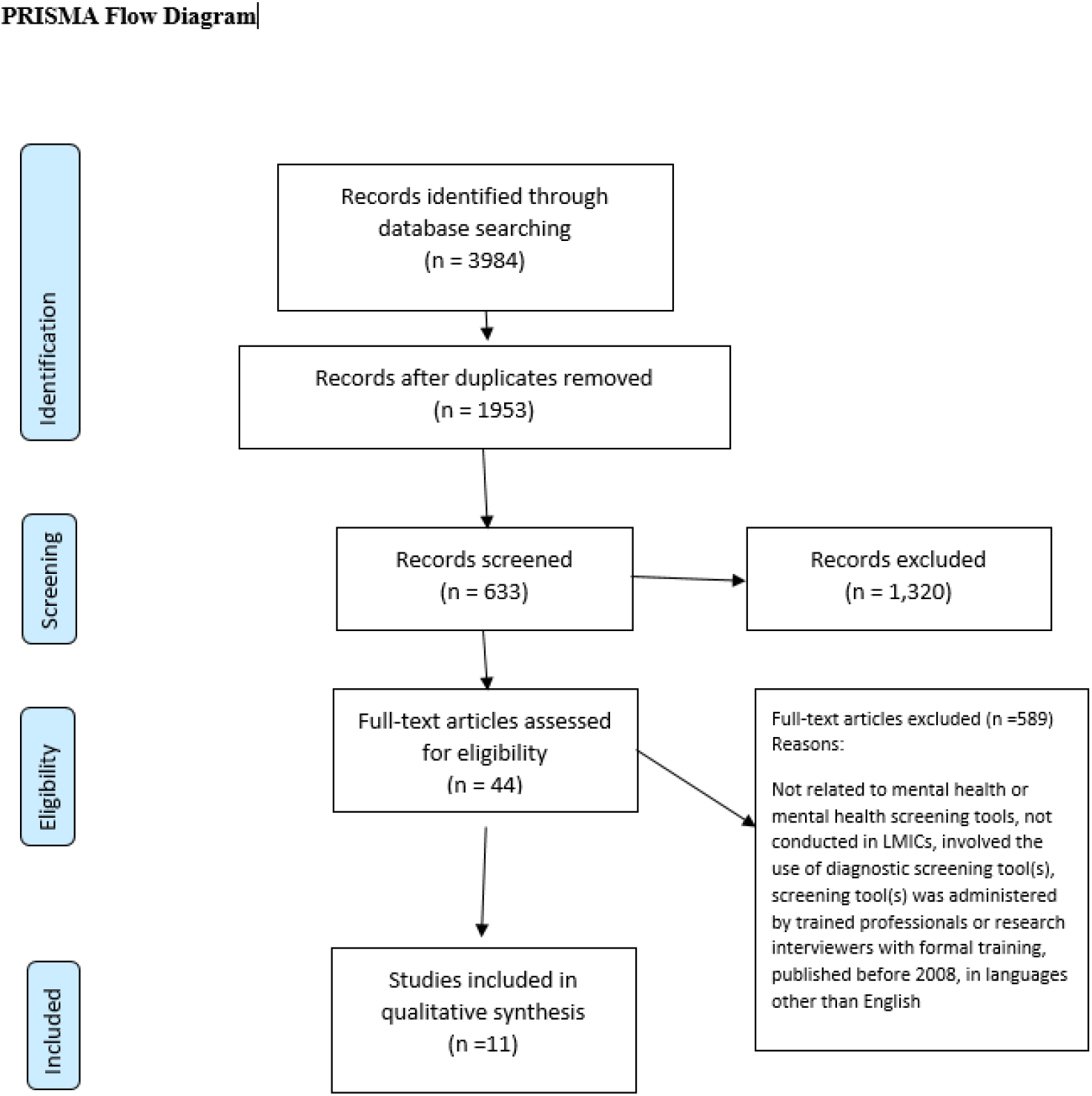
Study selection flow diagram.

### Study Characteristics

The majority of studies focused on the description or application of laypersons service provision rather than specific evaluation of screening tools. Although the primary focus of these studies was not on the use of screen tools per se, these studies were included as they provided a description of a number of screening tools using within lay person contexts. Four studies evaluated the effectiveness of community-based/primary health care level mental health interventions provided by lay community workers among different populations (20,21,24,25). Two of the studies describe community-based interventions provided by community lay workers trained to identify, counsel and refer people affected by disaster(22,23). These studies describe use of mental health screening tools by laypersons for screening, delivery or monitoring of MNS conditions in community-based mental health programs in LMICs. Within these studies, twelve screening tools were used. Only one study had the explicit purpose of evaluating mental health screening tools, a systematic review of mental health screening tools that have been validated for use in LMICs(14). This review included tools that were designed for use by lay-persons as well as other tools. We also searched the references of this review to determine if there were additional relevant studies to those identified in our review. We identified four additional studies that described the validation of several tools for use by lay persons in a community or PHC setting(26–29). Find the information in Annex II.

Across all studies, twelve screening tools were identified. Table 1 below provides a description of the identified mental health screening tools validated for layperson use in low and middle income countries.

**Table 1:** Summary table of the identified mental health screening tools validated for layperson use in low and middle income countries.

| **Screening tool** | **Definition of lay person** | **Length of tool(# of items)** | **Mode of Administration** | **Mental Health Condition(s)** |
| --- | --- | --- | --- | --- |
| Substance abuse and mental illness symptom screener (SAMISS) | Lay adherence counsellors (LAC). LAC had less previous knowledge of mental disorders and are of a lower professional category | 13/16 items | paper/computer-administered | multiple mental health and substance use conditions |
| International HIV Dementia Scale (IHDS) | Lay adherence counsellors (LAC). LAC had less previous knowledge of mental disorders and are of a lower professional category | 4 items | paper/computer-administered | Sub-cortical dementia associated with HIV |
| Shona Symptom Questionnaire | Lay workers are literate female elders supporting community health programs | 14 items | Paper | Multiple mental health conditions |
| Harvard Trauma Questionnaire (Section 4) | Lay community workers | 16 items | Paper | Posttraumatic stress disorder |
| Clinical Interview Schedule-Revised | Medical students (for this study) | 14 symptoms group questions | Computerized | Multiple mental health conditions |
| Aga Khan University Anxiety and Depression Scale (AKUADS) | Minimally trained adult females from the community that can read and write lingua franca Urdu | 25 items | Paper | Depression and anxiety |
| 12-Item General Health Questionnaire -12 (GHQ-12) | Lay people from the community trained over few days to conduct the assessment interview | 12 items | Paper | Multiple mental health conditions |
| 20-Item Self-reporting Questionnaire-20 (SRQ-20) | Lay people from the community trained over few days to conduct the assessment interview | 20 items | Paper | Multiple mental health conditions |
| Kessler Psychological Distress Scale (K-10, K-6) | Lay people from the community trained over few days to conduct the assessment interview | K-10 is ten-item. The K-6 score can be extracted from K-10 | paper/computer-administered | Multiple mental health conditions |
| 9-Item Primary Health Questionnaire (PHQ-9) | Lay people from the community trained over few days to conduct the assessment interview | 9 items |  | Multiple mental health conditions |
| 30-Item Geriatric Depression Scale (GDS-30) | lay-interviewers with at least 11 years of schooling were selected from the community | 30 items | Paper | Depression |
| Edinburgh Postnatal Depression Scale (EPDS) | female high school graduates (tenth grade and above) who had received two days of training in administration of the EPDS and SRQ-20 | 10 items | Paper | Multiple mental health conditions |

| Screening tool | Age range to which it applies | Setting(s) in which it has been used | Psychometric properties (Validity and reliability) | Country of study | Does use require training? |
| --- | --- | --- | --- | --- | --- |
| Substance abuse and mental illness symptom screener (SAMISS) | Adult | Clinic | specificity (58%), Sensitivity (94%), criterion validity (0.76) | South Africa | Yes |
| International HIV Dementia Scale (IHDS) | adult | Clinic (HIV specific setting) | specificity (79%), Sensitivity (45%), criterion validity (0.64) | South Africa | Yes |
| Shona Symptom Questionnaire | Adult | Clinic | specificity (83%), Sensitivity (67%), criterion validity (0.88) | Zimbabwe | Yes |
| Harvard Trauma Questionnaire (Section 4) | Adult | Community (Post-disaster) | specificity (73%), Sensitivity (87%), criterion validity (0.83) | Haiti | No. Self-administered, but questions were read out to those that cannot read There is manual for scoring |
| Clinical Interview Schedule-Revised | Adult/adolescent | clinical community | Specificity (96%), Sensitivity (100%) | Malaysia | Yes |
| Aga Khan University Anxiety and Depression Scale (AKUADS) | Adult | community | Specificity (81%), sensitivity (74%), | Pakistan | Yes |
| 12-Item General Health Questionnaire -12 (GHQ-12) | Adult, Older adults (>75 years) | Community, population-based study | Specificity (90%), Sensitivity (73%) and criterion validity (0.89). | India, Brazil | The interviewers received one week training |
| 20-Item Self-reporting Questionnaire-20 (SRQ-20) | Adult | Primary Health Care, community outreach | sensitivity of 85.7%, specificity of 75.6% and internal consistency of 0.84, | Ethiopia, India | The interviewers received one week training |
| Kessler Psychological Distress Scale (K-10, K-6) | Adult | Primary Health Care | Sensitivity of 65% and 58%, specificity of 89% and 91% and internal consistency of 0.8 and 0.74 respectively | India | The interviewers received one week training |
| 9-Item Primary Health Questionnaire (PHQ-9) | Adult | Primary Health Care | internal consistency of 0.79 | India | The interviewers received one week training |
| 30-Item Geriatric Depression Scale (GDS-30) | Older adults (>75 years) | Population-based study | Sensitivity of 73.3%, specificity of 65.4%, internal consistency of 0.87 and criterion validity of 0.74. | Brazil | Not specified |
| Edinburgh Postnatal Depression Scale (EPDS) | Adult | Community | Sensitivity of 76.5%, specificity of 36.1% and internal consistency of 0.47 | Ethiopia | Lay interviewers received two days of training |

### Analysis of the Identified Layperson Mental Health Screening Tools

Most of the layperson screening tools identified were originally developed and validated as self-report tools. Several have been administered by interviewers, especially in contexts where literacy levels of the target population is low(14). Table 3 provides a summary classification of the identified screening tools. Analysis of the identified tools shows that most tools were being used as interviewer-administered tools even though they were originally developed and validated for self-report. Information was not available as to whether this difference in mode of administration affects the measured outcome or score on these instruments.

For this study, we included mental health screening tools that have been developed or validated and used at the community level by lay persons that are non-mental health professionals and non-clinicians or non-health workers supporting mental health program or intervention (e.g. teachers, parents, peers, local/community workers). A systematic review of validated mental health screening tools in LMICs identified 21 screening tools that were validated for use by lay interviewers. However, all these tools were used by research assistants or trained interviewers rather than lay health workers in applied settings. For this current study, twelve layperson mental health screening tools that have been used in applied settings are described in Table 1. Some of these tools were developed in the western/high income countries and validated for use in LMICs while others were developed primarily for LMICs, including the SAMISSI, IHDS and AKUADS. Some were developed for specific mental health conditions (EPDS, GDS, HTQ and IHDS) while others are for common mental disorders. EPDS, GDS and IHDS were also developed for use in specific population ante- or post-natal women, geriatric and people living with HIV/AIDS respectively.

#### Substance abuse and mental illness symptom screener (SAMISS)

The SAMISS consists of 13-item or 16-items paper/computer administered screening tools developed to identify alcohol, substance use and common mental disorders (anxiety, depressive, adjustment and bipolar disorders) among people living with HIV/AIDS (PLWHA). The 16-item version contains items from alcohol use disorder identification test, 2-item conjoint screener, composite diagnostic interview and some items specifically designed for SAMISS. Initially developed in the USA, the SAMISS has been validated for use by lay counselors among people living with HIV/AIDS in South Africa with specificity (58%), sensitivity (94%) in detecting symptoms of common mental illness and substance abuse and significantly correlated (0.76) with the Mini International Neuropsychiatric Interview (MINI) administered by a mental health nurse(20).

#### International HIV Dementia Scale (IHDS)

The IHDS is a 4-item paper or computer administered screening tool specifically designed to assess sub-cortical dementia associated with HIV in different cultures, and by people with no formal training in neurology. The four items assess memory registration, motor speed, psychomotor speed and memory recall. The specificity and sensitivity of IHDS in detecting HIV dementia has been evaluated among in USA and Uganda(30). It has also been validated against neuropsychological test battery for use by lay technicians among PLWHA in South Africa with specificity (79%), Sensitivity (45%), criterion validity (0.64) in detecting subcortical dementia in this population(20). In the validation studies among South Africans living with HIV, researchers have noted that cultural, linguistic, or education of lay assessors may have affected the criterion validity of the IHDS. As such, further evaluation of the IHDS among lay assessors is warranted.

#### Shona Symptom Questionnaire

This tool was designed to provide indigenous and culturally-relevant mental health screening among indigenous populations in Sub-Saharan Africa. It is a 14-item common mental disorders (CMD) screening tool, adapted from the self-reporting questionnaire-20 (SRQ-20)(31), was developed for Shona speaking countries (Zimbabwe, Botswana and Mozambique) to assess common psychiatric symptoms and responses in binomial format. Five items are measures of indigenous idioms of distress of mental disorder that were not captured by SRQ-20. Its validity has been established among adolescent and young adult population in Zimbabwe by researchers(32). It was validated against SRQ-20 at optimal cut off point of five or more. Validation testing of the SSQ among the adult population shows good psychometric properties with specificity of (83%), Sensitivity (67%) and criterion validity (0.88) in detecting symptoms of depression and other CMD(32). It was used by lay worker (community health promoters) to screen PLWHA for symptoms of depression and other common mental disorders in Zimbabwe(21).

#### Harvard Trauma Questionnaire (Section 4)

This tool was used by lay health workers as a self-report instrument to monitor PTSD symptoms among survivors of 2010 Haiti earthquake, but questions were read out to those that cannot read (22). The original version of HTQ has four sections assessing history of traumatic event, personal description of the event, injury to the head and trauma symptoms. In this study only the first 16 items in section 4 were used to assess posttraumatic symptoms. Validity of the self-administered French version of this tool has also been established among survivors of torture and organized violence from sub-Saharan Africa(29). The content of the original version was first assessed for cultural relevance and adapted as appropriate. It was then translated into French using the Brisling’s back-translation method after which the French version was validated against Structured Clinical Interview for DSM (SCID). The validity study shows that HTQ is a good tool for assessing PTSS among the study population with specificity (73%), Sensitivity (87%) and criterion validity (0.83).

#### Clinical Interview Schedule-Revised

Although this was originally developed as a fully structured diagnostic instrument (Structured Clinical Interview Schedule-CIS) for use by psychiatrists, the modified/revised version had been developed to be used by trained lay interviewers or as self-administered questionnaire in assessing minor MNS conditions in the non-specialist settings. It is used to screen for the following 14 psychiatric symptom groups among adolescent and adults: (1) Somatic symptoms; (2) Fatigue; (3) Sleep problems; (4) Irritability; (5) Physical health worries; (6) Depression; (7) Depressive ideas; (8) Worry; (9) Anxiety; (10) Phobias; (11) Panic; (12) Compulsive behaviors; (13) Obsessive thoughts; (14) Forgetfulness/concentration problems. Scores on each symptom group ranged from .0 to 4 (and 0 to 5 for depressive ideas). The higher the score the higher the level of symptomatology. There are computerized self-administered version and interviewer administered versions suitable to be used by trained lay interviewers in assessing minor psychiatric morbidity in the community, primary health care and general hospital. The computer algorithm format of this tool enables generation of ICD-10 diagnosis without psychiatric consultation using the Programmable Questionnaire System (PROQSY). The Malay version of CIS-R has been validated against the structured clinical interview for DSM (SCID) ‘for use by lay interviewers among adult population in Malaysia which showed 100% sensitivity and 96 % specificity at a cut off score of 9.

#### Aga Khan University Anxiety and Depression Scale (AKUADS)

The AKUADS is designed to work as a self-administered or lay interviewer-administered depression and anxiety screening tool. The 25-items questionnaire made up of 13 psychological and 12 somatic items was developed from verbatim notes taken from persons speaking lingua franca Urdu to describe their symptoms of anxiety and depression(33). Each item has four response options scored from 0 – 3 with a cut-off score of 19 for positive screening test. It was validated for use by community health workers (CHW) against the psychiatrist's interview to detect depression and anxiety among adult population with specificity of 81% and sensitivity of 74% at cut off point of 19(34). It has also been used by trained lay community women to detect and monitor symptoms of anxiety and depression in new mothers receiving lay counseling(25).

#### 12-Item General Health Questionnaire

Is a 12-item self-reporting screening tool originally developed in the United Kingdom as a brief and general measure of psychiatric wellbeing assessing anxiety, depression, social dysfunction and loss of confidence(35). Each item assess the severity of a mental problem over the past few weeks using a 4-point Likert-type scale (from 0 to 3). The score was used to generate a total score ranging from 0 to 36. The positive items were corrected from 0 (always) to 3 (never) and the negative ones from 3 (always) to 0 (never). High scores indicate worse health. It has been validated for use by lay interviewers (lay community workers) against CIS-R as gold standard among adults in primary health care setting in India with specificity of 90%, Sensitivity of 73% and criterion validity of 0.89 in detecting symptoms of CMD (25,27).

#### Edinburgh Postnatal Depression Scale

Is a ten-item scale asking about common psychiatric symptoms experienced in the preceding week. Amharic version of EPDS was validated for use by lay interviewers (female high school graduates of tenth grade and above) against psychiatrist assessment using Comprehensive Psychopathological Rating Scale (CPRS) among post-natal women during a vaccination outreach in Ethiopia. It performed poorly in detecting CMD and MDD when compared to CPRS with internal consistency of 0.47, sensitivity of about 77% and specificity of 36%(26). One of the problems identified in the application of this tool among the study population was that it was difficult to translate some items to Amharic hence it was not well understood by the participants.

#### 20-Item Self-Reporting Questionnaire (SRQ)

The 20-item questionnaire has been evaluated in Ethiopia against the EPDS. The SRQ-20 had better psychometric properties in detecting CMD among the Ethiopian study population compared to EPDS, with internal consistency of 0.84, sensitivity of 85.7% and specificity of 75.6% when evaluated against CPRS(26). This is similar to the result from India where SRQ-20 shows internal consistency of 0.8 when compared against Revised Clinical Interview Schedule (CIS-R). This shows that SRQ-20 is a valid instrument to detect CMD among the study population when used by lay assessors.

#### 30-Item Geriatric Depression Scale

The Geriatric Depression Scale is a self-report 30-item questionnaire to assess depression in older people(36). Scores of 0-9 are considered normal, 10-19 indicate mild depression and 20-30 indicate severe depression. A validity study of the tool administered by lay-interviewers (with at least 11 years of schooling) who were selected from the community was conducted against the gold standard of the Schedules for Clinical Assessment in Neuropsychiatry (SCAN) among the general population age 75years and above in Brazil(28). The result shows sensitivity of 73.3%, specificity of 65.4%, internal consistency of 0.87 and criterion validity of 0.74 in detecting geriatric depression(28). This shows that 30-GDS is a valid instrument for use by lay assessors in detecting depression among the study population.

#### Kessler Psychological Distress Scale (K-10, K-6)

The K10 is a 10-item questionnaire developed to measure anxiety and depression. A shortened 6-item version of the questionnaire (K6) has also been advocated as a screening measure. There are self-report and interviewer-administered versions. The validation study of the use of K-10 and K-6 to detect CMD by lay interviewers against CIS-R among adults attending PHC in India shows sensitivity of 65% and 58%, specificity of 89% and 91% and internal consistency of 0.8 and 0.74 respectively.

#### 9-Item Primary Health Questionnaire

The nine-item PHQ (PHQ-9) is the depression screening module of the full PHQ, a self-administered version of the Primary Care Evaluation of Mental Disorders (PRIME-MD) diagnostic instrument for CMDs. It has been used for screening depression among primary-care patients. It is brief and has the ability to establish DSM-IV-based diagnosis of major depression. Its validity to detect CMD by lay interviewers among PHC attendees in India has also been established as poor internal consistency of 0.79. Sensitivity and specificity data were not reported in the reviewed study.

## Discussion

The lack of understanding of MNS issues and the huge treatment gap in LMICS is in part due to lack of information about the magnitude of MNS conditions, inability to understand the determinants of mental health issues and inadequate knowledge on how best to direct policy for improving support. This information is lacking because most of the mental health screening tools used in LMICs focused only on psychiatric symptoms in small sample survey or research. Incorporating the use of comprehensive screening tools used routinely or in large epidemiological studies will be able to provide comprehensive information that can inform integrated treatment planning at individual level and broader understanding of the needs and available resources in the community. Lay person health workers could play a key role within such initiatives if they are provided the right tools to accurately screen for mental health conditions.

There are a number of strengths of the lay person screening tools identified in this review. These include the relative brevity of most tools, the ease of administration among tools with bivariate responses, the minimal training requirements, low literacy requirements for completion, and the ability some tools to detect psychiatric conditions in physically ill patients. They also have strong psychometric properties in the study populations. In terms of limitations, most of these tools assess psychiatric symptoms alone and are restricted to the somatic manifestations usually ignoring the cognitive and emotional domains. Also, most of these tools were originally developed for use in the western world (except for SSQ and AKUADS) and translated into other languages to be used in other countries. Therefore care should be taken in interpreting the psychometric properties of the translated tools whose content might not necessarily be the appropriate cultural or indigenous idioms for mental distress in that population. Furthermore, the validation of some of these tools were done by health or mental health professionals in health facilities while the tools are expected to be used by laypersons for screening in the general population. Therefore there is need for more research exploring whether or not using a self-reporting tool as an interviewer-administered tools have any effect on the measured outcome or score. SSQ and AKUADS while they are culturally-relevant tools with good psychometric properties, are limited in their scope to assess comprehensive needs of the target population.

Khenti et al, 2015 proposed that mental health interventions, such as the development of a comprehensive mental health screening tool, should be developed by considering the five pillars, or the holistic policy framework, of global mental health. The five pillars are consideration of the broader determinants of mental health and sociocultural relevance of the mental health interventions. It consults relevant stakeholders especially, including the target population, in the design, development and monitoring of the intervention. Engagement of the target population provides opportunity for capacity building and reciprocal learning as well as sustainability of the intervention. Kentia et al recommended that when designing a mental health intervention, the knowledge or contribution/input from the people/community in which the tools will be used should be considered. For instance, in the case of the tools reviewed for this study there is little evidence that they have been developed with input from local communities. The tool should be adapted, validated and its reliability tested by taking into consideration the unique sociocultural belief of that particular context. In addition to using co-design with local communities, screening tools should assess not just MNS conditions but explore physical/medical, social and economic factors that can influence the mental wellbeing of the person. Furthermore the use of the tool should provide opportunity for knowledge acquisition either through training of the user/administrator or the information provided to the patient/caregiver during the screening process. Lastly the tool should be easy to access and use.

In Fig 3 we are proposing a framework for developing and choosing tools for use by lay persons to screen for mental health and addictions This reference framework will lay the foundation for designing or selecting an integrated mental health screening tool that can be used by laypersons with the general population at community or primary care level in LMICs or low resource settings.

**Fig 3:**
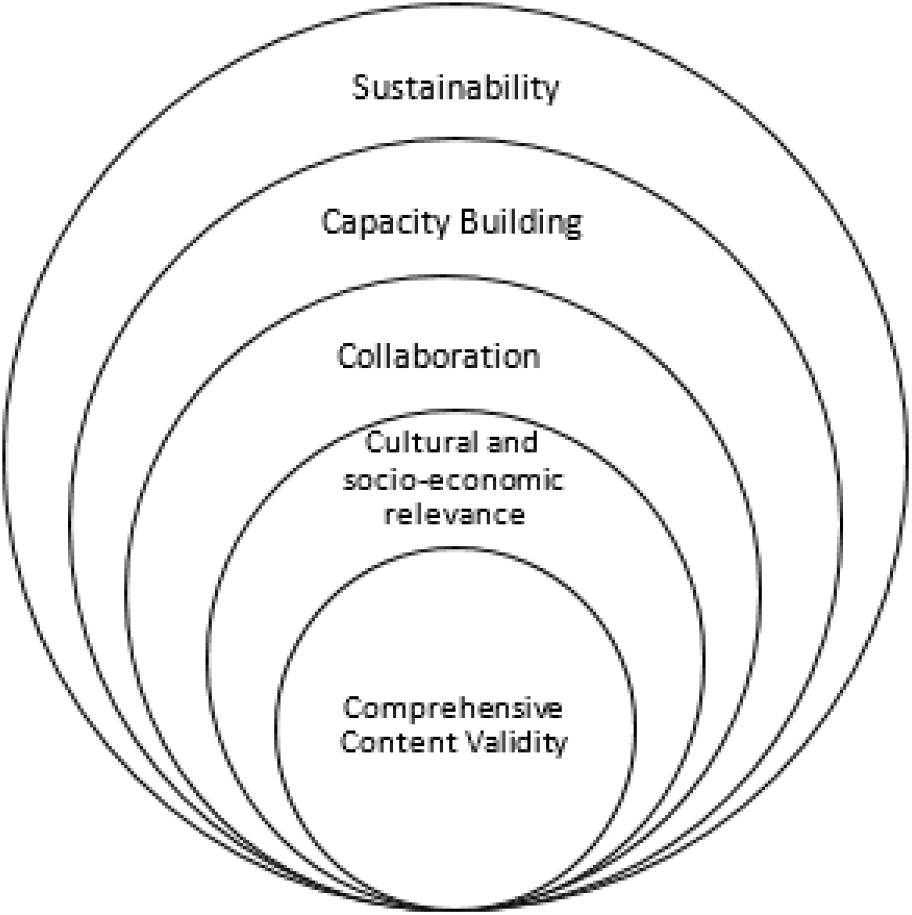
Five pillars of mental health screening tools (Adapted from the 5 Pillars of Global Mental Health and Addiction Work)

### 1. Comprehensive Content Validity

Validity is the extent to which a measurement method measures what it is intended or supposed to do, or the range of interpretations that can appropriately be placed on a measure(37). Validation studies can be done in clinical, research or community-based settings to evaluate whether the too is valid for the purpose it was developed and whether it was applicable in the particular context. Aspects of validity testing to consider could include content, criterion and construct validity. Content validity could be checked using the content matrix/table and having experts in the field review the technical content of the tool.

#### a) Technical Content

Technical content of a screening tool should be comprehensive in assessing psychiatric conditions, non-mental condition as well as other factors contributing to general wellbeing. In the reviewed studies some of the tools are used to screen for specific psychiatric conditions while some are used for general psychiatric symptoms. Some are designed for use in the general population while some are designed for use in a particular population. None of the screening tools were able to provide information about non-mental issues that might be contributing to the relevant mental condition.

The content of a comprehensive screening tool should contain the following groups of items:

i. Items that are common to all health issues at all level of care such as cognitive skills for decision making, communication, functional status, activities of daily living (e.g., personal hygiene, toilet use, eating), mood (e.g., negative statements, persistent anger, crying/tearfulness), behavior problems (e.g., verbal abuse, resisting care), falls, and physical/medical health symptoms (e.g., pain frequency and intensity, fatigue).
ii. Items common to social and other relevant services and or broader determinants of health such as instrumental activities of daily living (e.g., meal preparation, financial management, phone use), stamina, additional health conditions (e.g., extrapyramidal symptoms, abnormal thought processes, delusions), medication adherence, and preventive interventions and screening (e.g., influenza vaccination, breast screening), hearing aid use, social support and life events, family/close friends feeling overwhelmed by the person's illness, environmental factors
iii. Specialized items that are specific to mental health such as mental state indicators including number of lifetime psychiatric admissions, unrealistic fear or panic, intrusive thoughts or flashbacks, mood disturbance, command hallucinations, suicidal ideation, use of illicit drug, police intervention for criminal behavior, history of sexual violence or assault as perpetrator.

#### b) Psychometric properties

The criterion validity could be checked by comparing different domain of the screening tool with gold standard for each domain of the tool and construct validity could be checked by comparing the result of the screening tool in two extreme groups (e.g. those with and those without MNS conditions). Construct validity could also be evaluated using the convergent/divergent approach(37). Reliability testing measures how reproducible the results of the tool are under different conditions(37). Aspects of reliability testing to consider include internal consistency, inter-rater and intra-class reliability. Internal consistency which is calculated by Cronbach’s alpha measures how well the items in the measure correlate with each other to determine whether the items all seem to be measuring the same thing. If the mode of administration of the screening tool would be self-administered, the intraclass Correlation Coefficient (ICC) will be used for test-retest or intra-rater reliability in addition to internal consistency (Cronbach’s alpha). For interviewer-administered mode of administration, the inter-rater reliability (Interclass Correlation Coefficient) will also be checked. Pearson correlation coefficient could also be used but while ICC gives consideration to errors/biases that two raters might introduce into the measure, Pearson correlation has been found to be theoretically incorrect in this aspect(37). Percentage agreement is also commonly used, as is kappa and weighted kappa statistics. While ICC and Kappa yield identical results, ICC might be easier to calculate. There is also the alternate form reliability testing which requires creating another version of the tool although this is rarely used(37). Reliability of the screening tool could also be tested or piloted in an heterogeneous sample to evaluate the reliability of the tool to detect the defined attributes in people at risk and those not at risk of developing MNS condition. Evaluation of the psychometric properties of the tools should be done at similar settings in which the tools will be used. Since the screening tool will used among general population or those at risk of MNS conditions to make a decision whether an individual should be referred for further evaluation and possible treatment, the result of the reliability testing (Cronbach’s alpha, the intraclass or interclass correlation coefficient) higher than 0.7 might be considered good reliability of the tool in the target population compared to diagnostic tools that might require higher level of reliability. The ability of the tool to correctly identify those that are actually at risk of or with MNS condition (sensitivity) and those that are actually not at risk of or without MNS condition (specificity) could also be tested. Effort should be aimed at achieving high sensitivity of the measure to minimize false positive result due to high stigma associated with MNS condition in the target population.

#### c) Validation for layperson

Many validation studies for most of the layperson mental health screening tools in LMICs shows good psychometric properties. However, it is important to note that selection and use of screening tools developed in another context in a different cultural setting without proper validation can result in inaccurate results. Reliability and validity of instruments should consider the assessor and context. Many of the tools reviewed here were originally designed and validated either as self-report tools or as completed by a clinician or researcher; further validation in the lay person context was required. It will be beneficial to examine whether or not the settings in which the tools have been validated/used will affect the score or outcome measurement of the tools. Ethical approval by appropriate research ethic committee should be obtained for the validation study and the research team should comply with the “Do no harm” principles.

### 2. Cultural and socio-economic relevance

The design, development and psychometric evaluation of the screening tool should be done considering the cultural and socioeconomic context in which it will be used. For new tools or tools requiring adaptation, local people should be engaged throughout the development/adaptation process. Language used should be comprehensible by the target population. The design, format and presentation of the tools should be culturally acceptable. Samples of the expected users should be trained to administer the tools among target common population and setting in a culturally-sensitive manner. The users should be trained using the instruction manuals for the tools after which they will complete the assessment for selected individuals from the general population. Information about the experiences of the users of the tools can be collected through focus group discussion or questionnaire. This approach will provide the opportunity to conduct real world assessment and training/capacity building of laypeople that will use the tools. Lessons learned, observations, feedback and recommendations from users and participants can be applied when improving the version of the tool.

### 3. Collaboration

The development of the tool’s content should be a collaborative effort of external mental health specialists, researchers and lay community members. The tool should be design such that it can be integrated with common clinical assessment. For instance the output or results for the layperson screening tool could provide basic understanding of the person’s needs, while the clinical assessment goes into greater depth to understand those needs in relation to treatment options. While specialists provide the basis for the technical contents of the tools and ability of the results to inform further intervention, the knowledge of the local people of the community where the tools will be used are important to ensure cultural relevance and acceptability. Collaboration in the development of user manuals is also important for maintaining the reliability and validity of the content by providing item descriptions, process instructions and examples. There should be communication and collaboration between community-lay person and the PHC or hospital staff for the use of the tool in practice. This is important to enhance supervision and opportunity for incremental training and support which can lead to *task shifting*.

### 4. Capacity Building (Training)

The development and the use of mental health screening tools should provide opportunities for training lay community workers that will administer these tools. In addition, the use of the tools with the general population or people affected by MNS conditions should provide information on awareness and improve their knowledge of MNS issues.

Self-report tools usually include instructions or come with separate instruction manuals for completing and scoring them and so does not require training. In the reviewed studies interviewers were trained on how to administer the tools and score the responses. Mental health screening tools should have accompanying instruction manual on how to complete and score them. These manual can be used for training the users, especially the computer-based algorithm type.

The assessors should be trained to use various information sources such as observation, interview with the person and those accompanying them (friends/family).

### 5. Sustainability

Sustainability will be the outcome of a tool that has been developed with adequate consideration of the initial four pillars. These pillars should not be considered as isolated pillars but all inclusive. The development of a holistic screening tool for a mental health intervention that is sustainable requires that the tool is accepted and demand for use which will depend on the ease of use (length of the tools, language used and cost). And finally the tool should be easy to adapt to different culture or setting.

*Acceptability and Utility:* One way to ensure acceptability and utility of a layperson mental health screening is to engage the community from the design/development of the tools through its validation, provide feedback on its use, contribution to evaluation of its effectiveness and review. This will promote community ownership. Also the use of the tool should provide interpretation of results or score and the next step to take. The data should be collected such that it can be easily integrated into the health information system.
*Ease of Use:* The screening tools are either in paper form, electronic form or computerized applications with each item coded. When electronic format is not available, paper form can be completed and records entered locally into the database. Each item should be in simple clear sentence(s) that can be easily understood. Each item should have standardized set of simple responses with clear definition and timeframe. While lay assessors would be trained on how to use all sources of information in completing the tools, this should not include clinical judgment although a clinician can also administer the tool. Also, while many of the short versions of the mental health screening tools focus on psychiatric symptoms only, the length of some of the majority of the tools reviewed depends on the version being used. Some tools have both long and short versions. The number of items on a screening tool should be appropriate for collecting adequate relevant information. Tools should be in the local language of the community which can be done through translation and back translation. People should be screened free of charge therefore the tool should be licensed for free and easily accessible.

The incorporation of the five pillars into the design of layperson screening tools is expected to improve the reach, utility, and impact of the lay person screening process supporting a more responsive health system. Therefore, in order to reach more people and create a more responsive mental health system, the effort needs to include developing a holistic tool that is of cultural relevance. This as a matter of fact cannot be achieved by a handful of people. Rather there is need to partner with different stakeholder or partners for knowledge exchange, and pool different innovations and experiences that these diverse ideas can bring. Capacity building is also not just focused on the researchers but also on the users of the tools. The use of the tools should create opportunities for creating or raising awareness about mental issues, increase the knowledge of the participants and their caregivers and help them in the understanding of the determinants of mental health. The interviewer should also use different sources to gather as much information as possible. There should be room for interaction between the clients and the assessor. There should be opportunities to ask questions and also receive feedback. And the feedback should be used to improve the design or the delivery of the tools.

### Limitations of the Study

This study did not do technical review and analysis of pooled psychometric data of the identified tools because this is not the purpose of this review. The purpose of this study is to conduct a qualitative review and examine the characteristics of mental health screening tools proposed for use by layperson as described above. This study also did not review grey literature reports which could also have biased the selection of the studies chosen for these review.

## Conclusion

The needs of people living with MNS conditions are multifaceted and interlinked in a complex manner. Inability to accurately identify these needs is a major contributor to the treatment gap in their management. Screening tools can provide comprehensive information about these needs to inform holistic care and responsive health system. Community layperson can reach more people in needs with information and access to care. Developing a layperson screening tools is vital for integrated community-based mental health intervention. This study has proposed a holistic framework that can be consulted for developing or selecting a layperson mental health screening instrument.

More research are needed to evaluate the practical application of this framework. Other research questions unanswered by this study include whether or not there are impact or effect on measured /outcome if self-report tools are used as interviewer-administered tool against being used as self-administered tool in order to know which version is more effective in low resource settings.

